# Amplification of frog calls by leaf substrates: implications for terrestrial and arboreal species

**DOI:** 10.1101/2020.11.02.361840

**Authors:** Matías I. Muñoz, Wouter Halfwerk

## Abstract

Signal detection is a minimum requirement for any communicative interaction. Acoustic signals, however, often experience amplitude losses during their transmission through the environment, reducing their detection range. Displaying from sites that increase the amplitude of the sound produced, such as cavities or some reflective surfaces, can improve the detectability of signals by distant receivers. Understanding how display sites influence sound production is, however, far from understood. We measured the effect of leaf calling sites on the calls of an arboreal (*Hyalinobatrachium fleischmanni*) and a leaf-litter specialist (*Silverstoneia flotator*) frog species. We collected the leaves where males of both species were observed calling, and conducted playback experiments to measure their effect on the amplitude of frog calls. Overall, the leaves used by *H. fleischmanni* and *S. flotator* were of similar dimensions, and amplified the calls of each species by about 5.0 and 2.5 dB, respectively. The degree of call amplification was unrelated to leaf dimensions or the position of the frogs on the leaves, but explained by the different frequency content of the calls of each species. Depending on the spatial location of intended and unintended receivers, we suggest that amplification of frog calls by leaves could represent either a benefit or impose costs for arboreal and terrestrial species. We argue that the microhabitat of the substrate from which animals display needs to be considered when addressing signal evolution.

**Lay summary:** Animals produce signals from specific locations in the environment, yet we know surprisingly little about the effects of the small-scale habitat on animal communication. Here we show that the calls of a terrestrial and an arboreal frog species are amplified by the leaves they use as calling sites. We argue that the consequences of this enhancement need to be considered in relation to the spatial location of intended (males and females) and unintended receivers (predators and parasites).

## Introduction

In animals, communication by means of sounds is widespread and crucially involved in activities such as mate choice, rival competition and parent-offspring interactions. Communication comprises three interacting components: a sender that produces the signal, the environment through which it propagates, and a receiver that perceives the sound (Bradbury and Vehrencamp 2011). Under this framework, the transmission environment is generally considered to impose constraints to communication. Propagating signals will experience a number of changes, including the modification of their acoustic structure (i.e., changes in their temporal and spectral attributes) and the loss of amplitude (Morton 1975, Wiley and Richards 1978, Ryan and Kime 2003). Spectro-temporal alterations can jeopardize the ability of receivers to recognize the signal and respond accordingly (Slabbekoorn 2004), but the loss of amplitude can have even more profound consequences for communication. This is because the ability of receivers to detect a signal is tightly linked to its amplitude, and signal detection is a fundamental step in any communicative interaction. If receivers fail to detect the presence of a signal, then communication will be completely hindered irrespective of other possible alterations in the sound structure.

The range over which a signal can be detected is mainly determined by the interplay of four variables: the amplitude of the signal at the source, the degree of environmental attenuation, the noise level at the receiver’s position, and the hearing sensitivity of the receiver (Marten and Marler 1977, Brumm and Slabbekoorn 2005). Environmental attenuation of acoustic signals is a well-described phenomenon. The amplitude of a sound decreases with increasing distance from the source, even in free-field conditions where there are not objects between the sender and receiver (Wiley and Richards 1978, Forrest 1994). However, the presence of sound reflective and absorptive elements (e.g., vegetation) is a typical feature of natural environments. These objects will induce additional amplitude losses to propagating sound, especially in the high frequency range (Wiley and Richards 1978). Additionally, sounds produced close to the ground often experience particularly high attenuation rates (Wiley and Richards 1978, Arak and Eiriksson 1992, Mathevon et al. 1996, Schwartz et al. 2016). Animals could, thus, reduce signal attenuation and improve detection distances simply by producing louder sounds, delivering them at lower frequencies, or from elevated display sites.

Louder signals are not only more likely to be detected, but can also influence mate choice. Acoustic signals presented at higher amplitude are preferred by females in a number of taxa (e.g., Gerhardt 1987, McKibben and Bass 1998, Ritschard et al. 2010). Importantly, females are able to discriminate relatively small amplitude differences. For example, females of many frog species are estimated to discriminate amplitude differences of 2-5 dB between two signals (see Bee et al. 2012 for a summary). However, high amplitude signals can also be detected and preferred by unintended receivers (e.g., Tuttle and Ryan 1981, Gomes et al. 2017). Acoustically-guided predators and parasites, will impose costs on the senders, and thus are expected to counter balance some of the mating benefits of high amplitude signaling.

While habitat-dependent attenuation can limit signal detection, other environmental features can enhance sound broadcast. In some species, signalers can use, build, or modify display sites that result in louder signals. For example, the burrows occupied by some frogs and crickets enhance the amplitude of their songs (e.g., Bennet-Clark 1987, Muñoz and Penna 2016). While these cavities seem rather complex display sites, other apparently simpler structures have the potential to generate similar effects. The surface of leaves, for example, are used by many insects and frogs for sound broadcast and could act as sound reflectors. Indeed, recent studies show that leaves have relevant acoustic properties across a surprisingly wide range of ecological contexts. Singing tree-crickets, for example, produce louder songs by creating so called baffles on leaves (Mhatre et al. 2017), whereas gleaning bats rely on ultrasonic reflections to detect motionless prey resting on leaves (Geipel et al. 2019).

Male frogs advertise their presence to potential mates and rivals using vocalizations. Different species broadcast these vocalizations from specific substrates, such as from the water, cavities, or the branches and leaves of trees. Here, we studied the effect of leaf calling sites on the call amplitude of two tropical species: a glass frog (*Hyalinobatrachium fleischmanni*) and a rocket frog (*Silverstoneia flotator*). *Hyalinobatrachium fleischmanni* belongs to the family Centrolenidae, and males vocalize during the night from leaves on trees and bushes next to streams (Fig. 1A). Males can either call on the top or the underside of leaves (Delia et al. 2010). On the other hand, *S. flotator* belongs to the family Dendrobatidae, and males call during the morning and afternoon while sitting on top of leaves in the leaf litter (Fig. 1B). Besides different activity periods and calling heights, males of both species share a number of reproductive behaviors, like the defense of territories where calling, mating, oviposition and paternal care take place (Savage 2002). The males of these frog species often come back to the same calling leaves during consecutive days, and thus are suitable candidates to study the acoustic consequences of calling from leaves.

**Figure 1:**
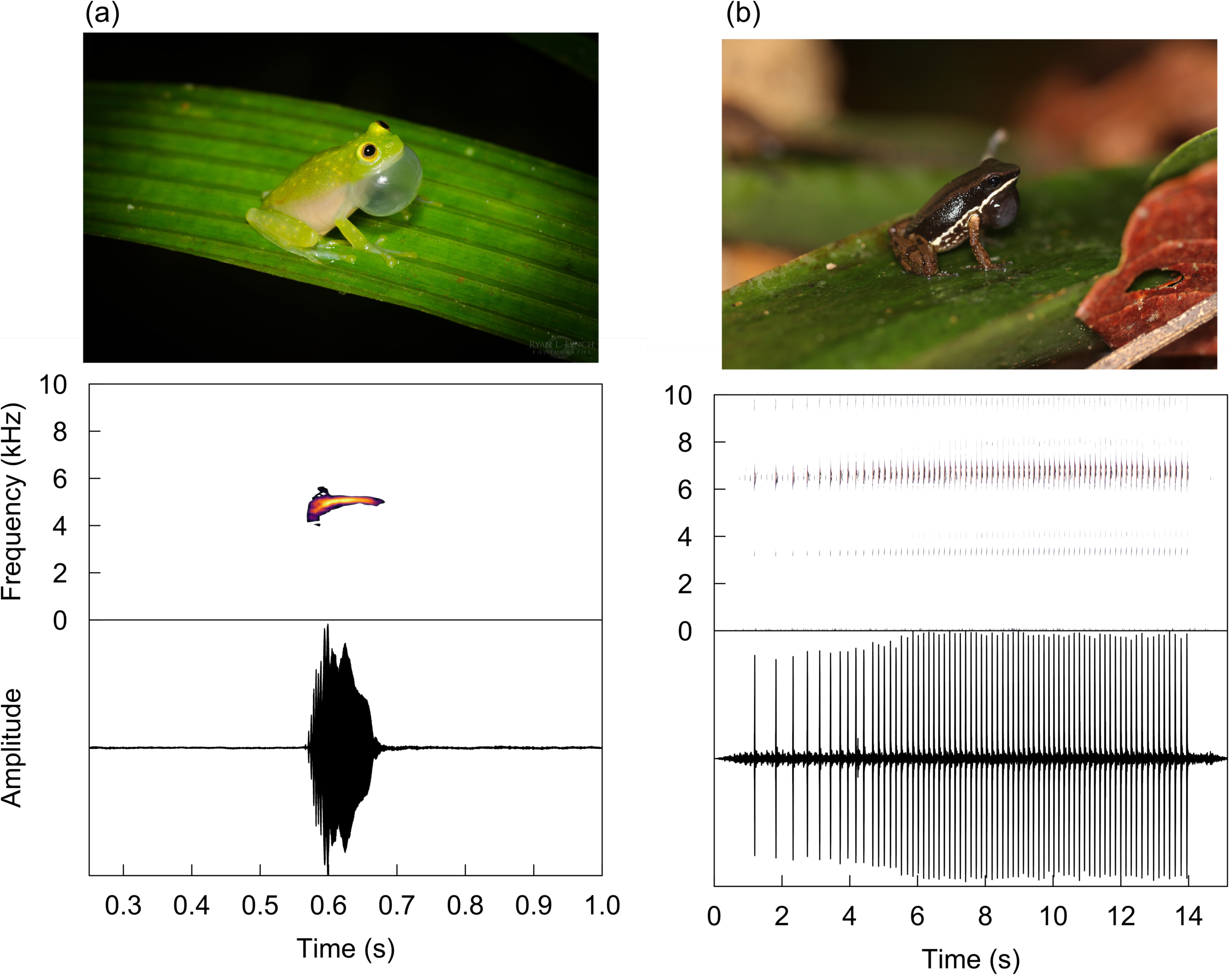
Calling male (a) *H. fleischmanni* and (b) *S. flotator* (top). Spectrograms and oscillograms (bottom) of an advertisement call of (a) *H. fleischmanni* and (b) *S. flotator*. Photos courtesy of Ryan Lynch and Rhett Butler.

## Methods

### Study area and leaf collection

The territories of male *H. fleischmanni* (N = 25) and *S. flotator* (N = 22) were surveyed between June and July 2019 in Barro Colorado Island (BCI), Panamá. Males were photographed at their calling sites and placed on a nearby leaf. Then, we collected the leaves used for calling and noted the exact position of males using a permanent marker. Leaves were transported to the facilities of the Smithsonian Tropical Research Institute in BCI for measurements. To reduce possible changes in the shape of leaves or other relevant properties, all the measurements were done within a few hours after collection.

### Leaf dimensions and frog calling position

Each leaf was photographed next to a reference scale, and using ImageJ (Schneider et al. 2012) we measured three variables related to their dimensions: maximum leaf width (cm), maximum leaf length (cm), and leaf area (cm^2^). First, we rotated the photographs to vertically align the apex and base of the leaf. The maximum widths and lengths were obtained from measuring the sides of a rectangle encompassing the apex, the base, and both sides of the rotated leaf image. The center of the leaf was defined as the geometric center of this rectangle. To measure leaf area, photographs were transformed to black and white and the color threshold adjusted to until the leaf area was discriminated from the background. Finally, we also measured two variables related to the position of the calling males on the leaves: distance to leaf center (cm), and distance to nearest leaf edge (cm). These two measures were obtained by drawing a circle centered on the frog position and measuring its radius until reaching the center and the nearest edge of the leaf.

### Acoustic stimuli creation

We assessed the acoustic properties of leaves using pure tones and natural advertisement calls. A sequence of 0.25s duration tones from 2.0 to 15.0 kHz in 0.5 kHz steps was created using Audacity 2.2.2. The relative amplitude of each tone was equalized to correct for the frequency response of the speaker used for the playbacks.

Advertisement calls of *H. fleischmanni* (N = 10 individuals) and *S. flotator* (N = 10 individuals) were recorded along the trails and streams of BCI. Vocalizations were recorded at a distance of 50 cm from either the front or side of the males using a microphone (G.R.A.S. 1/2 inch 46AE) connected to a digital recorder (Zoom H6). For each recorded *H. fleischmanni* individual we randomly selected 10 calls which were included in the audio file used for acoustic testing (Fig. 1A). Because the call of *S. flotator* consists of long sequences of rapidly repeated short pulses, a single vocalization of each recorded individual was included in the audio file (Fig. 1B). The peak frequency of *H. fleischmanni* and *S. flotator* calls was (mean±SD) 5.19±0.16 and 6.46±0.33, respectively.

### Acoustic properties of leaves

To measure the acoustic properties of leaves we employed a disk-shaped piezoelectric speaker (diameter: 35 mm, KEPO FT-35T-2.6A1-475) connected to a mini audio amplified (PAM8610) and a laptop computer (Apple Inc.). The piezoelectric was mounted on a thin vertical wire and placed at 50 cm from a microphone (G.R.A.S. ½ inch 46AE) connected to a digital recorder (Zoom H6) (Fig. 2). We covered the wall behind the piezoelectric, and the ground path between the microphone and the piezoelectric with acoustic absorbing foam to reduce unwanted sound reflections.

**Figure 2:**
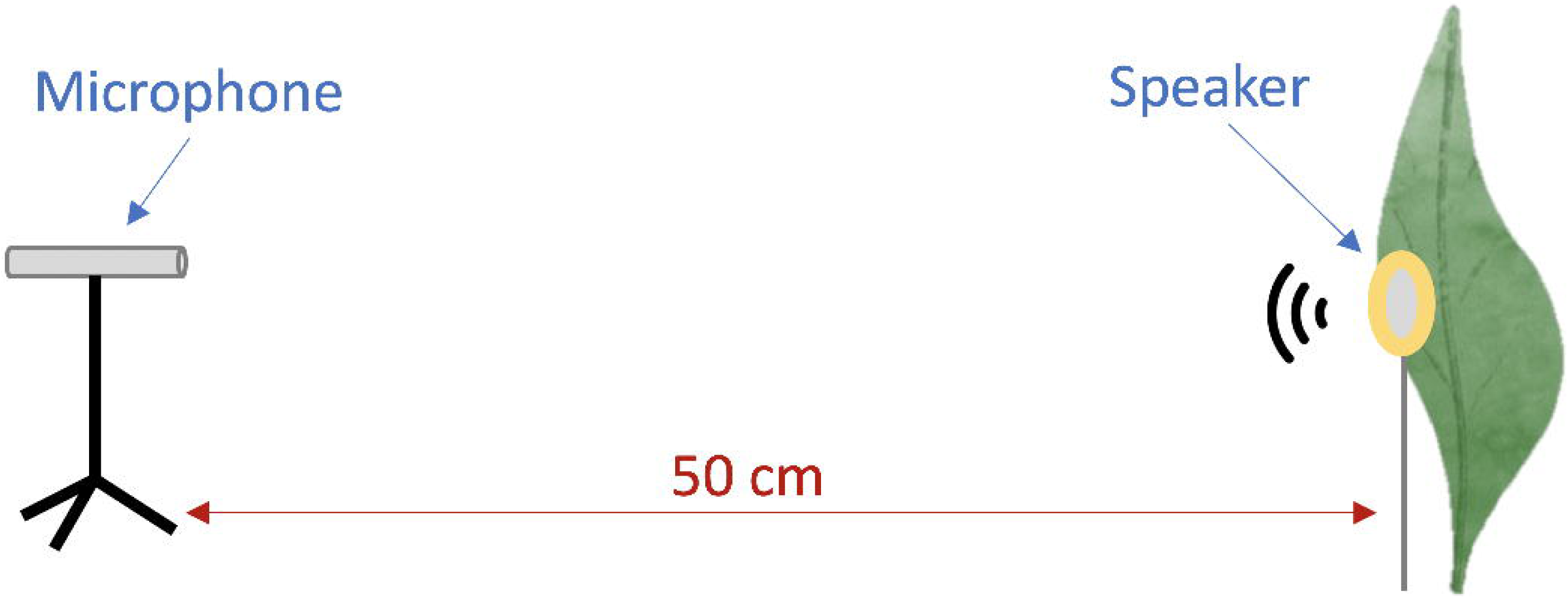
Experimental set up used to test the acoustic properties of leaves.

In each trial, a leaf was placed behind the speaker at a distance of about 0.5 cm from its back surface (Fig. 2), mimicking the position of a frog sitting on the leaf surface. The piezoelectric was centered at the exact position where the frog was observed calling, and care was taken not to alter the position of the piezoelectric relative to the microphone while placing and removing the leaf. Pure tones and frog calls were broadcast with the leaf in this position and recorded with the microphone. To compare the amplitude of tones and calls in the absence of the leaf, we obtained a baseline recording of these sounds before placing and also after removing the leaves from the set up. Leaves were tested only with the calls of the species observed calling on them, and the leaves used by *H. fleischmanni* were tested on the same side (upper or underside) where males were observed calling.

### Acoustic analyses

We measured the root-mean-square (RMS) amplitude of tones and calls using RavenPro 1.5 (Center for Conservation Bioacoustics 2014). For tones, recordings were band-stop filtered between 0 and 1.0 kHz to reduce the presence of low frequency sounds, and a 0.15 seconds segment from the middle of each tone was selected for RMS amplitude measurements. For the *H. fleischmanni* and *S. flotator* calls we measured the RMS amplitude in the 4.0 - 6.0 kHz and 5.0 - 8.0 kHz frequency bands, respectively. For each leaf we computed the amplitude gain of tones and calls using the following equation: Amplitude gain (dB) = 20*log_10_(RMS_leaf_/RMS_no leaf_). Where RMS_leaf_ corresponds to RMS amplitude measurements obtained with the tested leaf placed behind the piezoelectric buzzer, and RMS_no leaf_ corresponds to RMS amplitude values measured before placing the leaf. Positive amplitude gains indicate amplification due to the presence of the leaf, while negative values indicate attenuation. The variation between the two baseline measurements was minimal, and thus we only report results obtained using the values obtained at the beginning of each trial.

### Statistical analyses

All the statistical analyses were performed in R (version 3.6.3, R Core Team 2020). The five measures of leaf dimensions and frog position were compared between the two studied species using unpaired two-sample t-tests, or Wilcoxon rank sum tests if the normality assumption was not met. Additionally, to reduce the number of leaf dimension and frog position variables for further analyses, we used principal component analysis on the correlation matrix using the base function *prcomp()*. The five leaf measures were log_10_-transformed, centered and scaled for this analysis. Only principal components with eigenvalues > 1 were considered.

The amplification of pure tones was analyzed using generalized additive models (GAM) with the library ‘mgcv’ (version 1.8-33, Woods 2011). We fitted a GAM model with gaussian distribution and identity link function. We included the amplitude gain (dB) as dependent variable and the tone frequency (kHz) as a by-species smooth predictor. Leaf identity was included as a random smooth spline in the models to account for the repeated measures obtained on a same leaf. Model estimates and 95% confidence intervals were plotted using the library ‘mgcViz’ (version 0.1.6, Fasiolo et al. 2018), and used to visually evaluate significant amplification or attenuation of specific frequencies. Additionally, to compare the profile of tone amplification between both species we used spectral cross-correlation implemented in the R package ‘seewave’ (version 2.1.4, Sueur et al. 2008). Spectral cross-correlation is often used to compare the frequency spectra of two sounds, and here we used it to compare the mean tone amplification profiles of glass frog and rocket frog leaves. Cross-correlation coefficients go from -1 to +1, and are interpreted in a similar way as regular correlation coefficient. A cross-correlation coefficient of +1 indicates that the two spectra are identical, while a coefficient of -1 indicates they are completely inverse.

To evaluate whether the amplitude gains of advertisement calls differed from 0 dB (i.e., no effect of leaves on call amplitude) we used one-sample t-tests, or one-sample Wilcoxon signed rank test if normality assumption was not met.

To evaluate differences in call amplification between both species we used two-sample Wilcoxon rank sum test. Male *H. fleischmanni* called from either the top or the underside of leaves. We compared the amplitude gains of leaves where males called on top and the underside using two-sample Wilcoxon rank sum test.

We evaluated associations between the leaf dimensions and call amplification levels using ANCOVA. We fitted two ANCOVA models, one for each principal component obtained from the leaf dimensions analysis. In both models we included amplitude gain as the dependent variable, and the species and principal component (either PC1 or PC2) as independent variables. We initially included the statistical interaction between species and principal component as independent variable, but these terms were not significant and were dropped from the final models reported. For all the linear models fitted (GAMs and ANCOVAs) we visually inspected the residuals to evaluate deviations from normality.

Mean and standard deviation amplitude gain values of tones and calls were obtained using the *meandB()* and *sddB()* functions in the R package ‘seewave’ (version 2.1.4, Sueur et al. 2008).

## Results

### Leaf dimensions and frog calling positions

Male *H. flotator* called from leaves with a larger area and wider than male *H. fleischmanni* (Table 1). In contrast, leaves used by both species had similar lengths, and males of both species were equally distanced from the center of leaves (Table 1). Male *S. flotator* called further away from the edges of leaves relative to male *H. fleischmanni* (Table 1).

**Table 1:** Dimensions of the leaves used male *H. fleischmanni* (N = 25) and *S. flotator* (N = 22) for calling. Results of statistical tests used to compare the dimensions of leaves used by both species are also shown. Values in the table correspond to mean ± SD.

|  | *H. fleischmanni* | *S. flotator* | Statistic (d.f.) | P |
| --- | --- | --- | --- | --- |
| Maximum leaf width (cm) | 8.36 ± 3.60 | 9.75 ± 2.18 | W = 173.5<sup>†</sup> | **0.031** |
| Maximum leaf length (cm) | 17.72 ± 6.45 | 20.51 ± 5.44 | t = -1.59 (45)<sup>*</sup> | 0.119 |
| Leaf area (cm<sup>2</sup>) | 107.96 ± 87.25 | 140.85 ± 64.86 | W = 149<sup>†</sup> | **0.007** |
| Frog distance to leaf center (cm) | 4.57 ± 3.05 | 3.97 ± 2.87 | W = 315.5<sup>†</sup> | 0.394 |
| Frog distance to leaf edge (cm) | 1.49 ± 0.52 | 2.62 ± 1.18 | W = 133<sup>†</sup> | **0.003** |

<sup>\*</sup>Unpaired two-sample t-test; <sup>†</sup>Two-sample Wilcoxon Rank Sum test. Significant differences are shown in bold.

The first two principal components explained 81.5% of the variation in leaf dimensions and frog calling position, and were the only components with eigenvalues > 1 (Fig. 3, Table 2). The PC1 was inversely correlated with leaf width, length, area, and the distance of the frog to the center of the leaves (Table 2). The PC2 is directly correlated with the distance of males to the edge, and moderately and inversely correlated with the distance of males to the center of the leaves (Table 2). Overall, there was large overlap between the PC scores of both species, although glass frogs calling from small leaves (i.e., large PC1 scores) tended to do it closer to the leaf edges (i.e., small PC2 scores) as compared to the rocket frogs (Fig. 3).

**Table 2:** Principal component analysis of leaf dimensions and frog calling position measured for *H. fleischmanni* and *S. flotator*.

|  | **PC1** | **PC2** |
| --- | --- | --- |
| Maximum leaf width (cm) | **-0.552** | 0.118 |
| Maximum leaf length (cm) | **-0.414** | 0.212 |
| Leaf area (cm<sup>2</sup>) | **-0.570** | 0.122 |
| Frog distance to leaf center (cm) | **-0.446** | **-0.483** |
| Frog distance to leaf edge (cm) | 0.008 | **0.832** |
| Eigenvalue | 2.778 | 1.306 |
| Proportion of variance explained | 0.554 | 0.261 |

Interpretable factor loadings are shown in bold.

**Figure 3:**
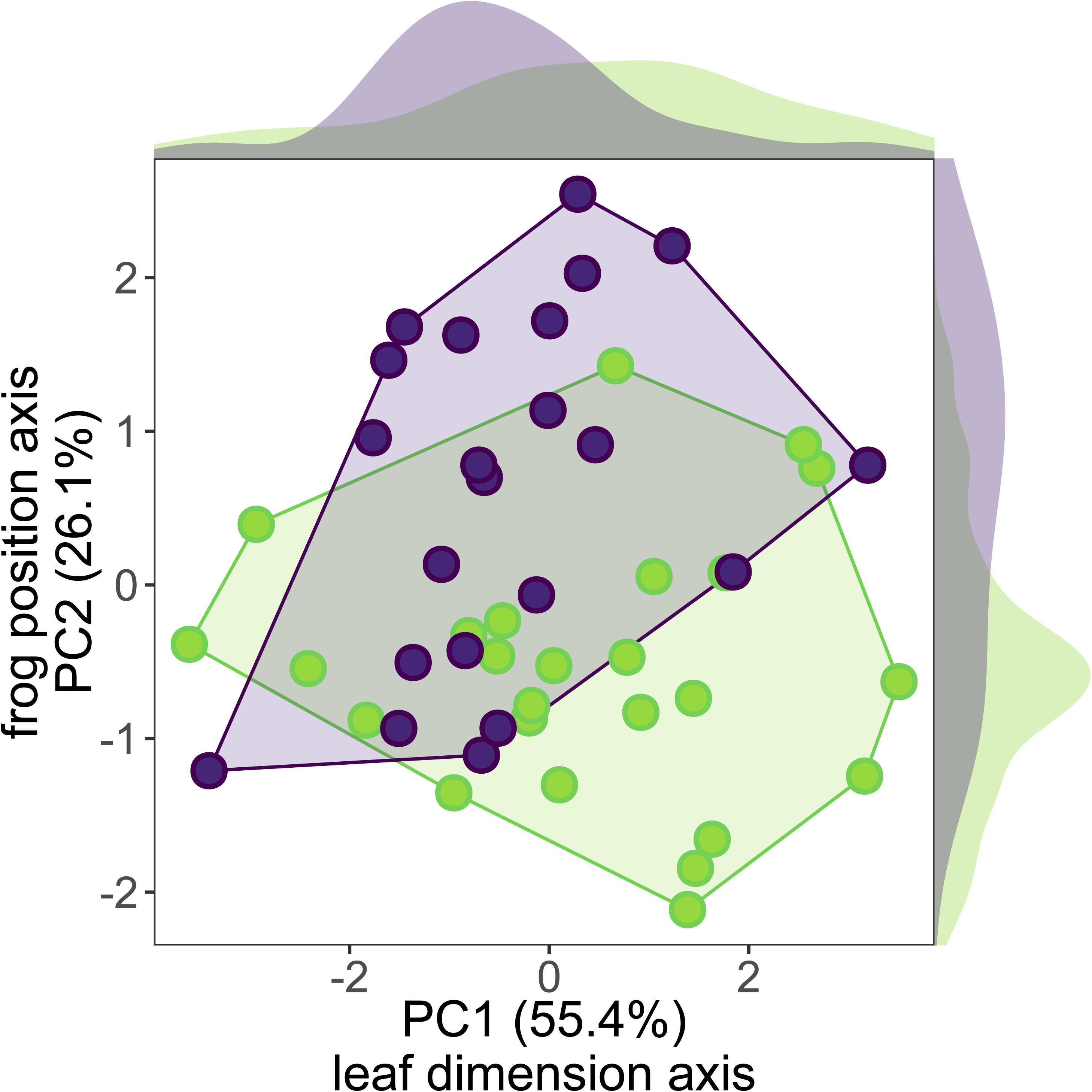
Principal component analysis of leaf dimensions and frog calling positions of *H. fleischmanni* (in green) and *S. flotator* (in purple). Distribution density plots for each variable are show on the outer margins.

### Amplification of pure tones

For both species, tones in the 4.0 – 6.0 kHz frequency range experienced larger amplification than other frequencies, and deviated from 0 dB as judged by the estimated 95% confidence intervals. This frequency band overlaps with the peak frequency range of *H. fleischmanni* calls (Fig. 4A), but not the calls of *S. flotator* (Fig. 4B). The leaves of both species showed similar pure tone amplification profiles, as indicated by a large and significant spectral cross-correlation coefficient between them (Spearman’s rho = 0.748, P < 0.001, frequency offset = 0 kHz).

**Figure 4:**
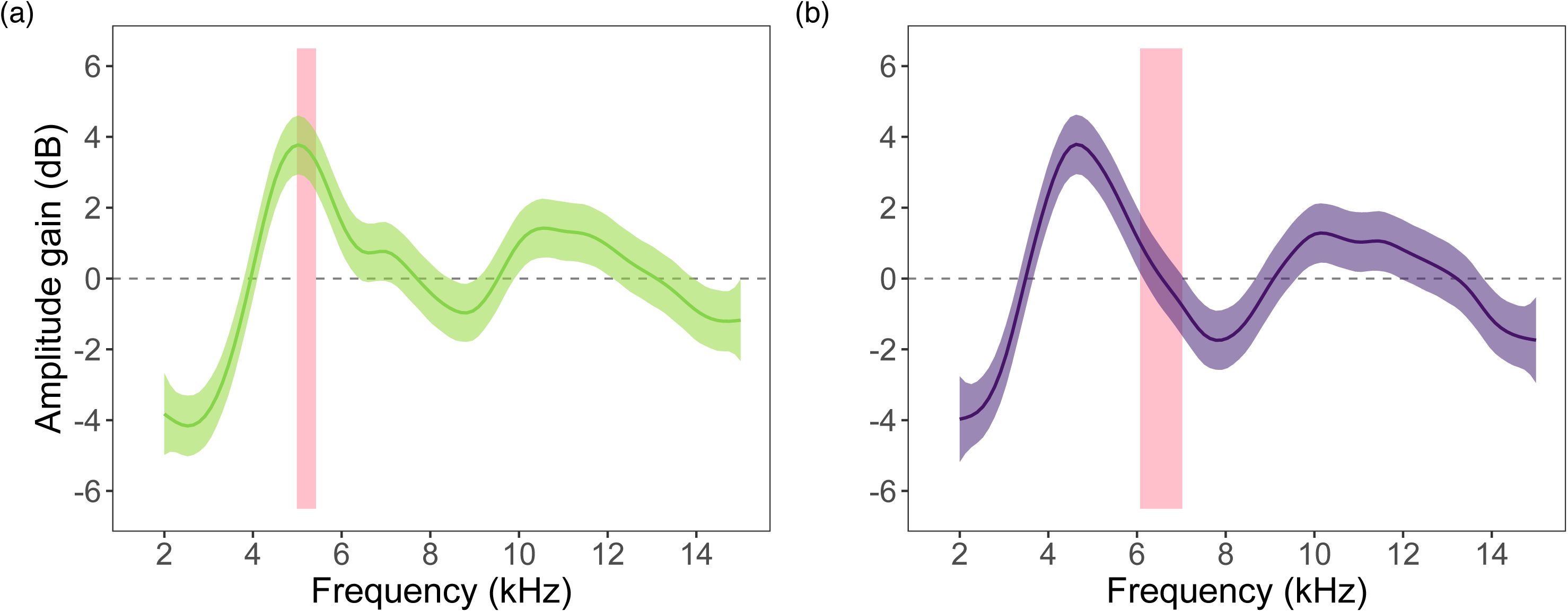
Amplitude gain of tones measured for leaves used by (a) *H. fleischmanni* and (b) *S. flotator*. Solid lines and shaded areas represent fitted amplitude gains and 95% confidence intervals estimated from the generalized additive model. The red vertical bars represent the range of call peak frequencies of each species.

### Amplification of calls

The calls of *H. fleischmanni* (N = 100 calls from 10 individuals) were amplified by (mean±SD) 4.93 ±1.39 dB, and the calls of *S. flotator* (N = 10 calls from 10 individuals) by 2.44±1.53 dB. For both species, call amplitude gains significantly deviated from 0 dB (*H. fleischmanni:* one-sample Wilcoxon signed rank test, W = 324, P < 0.001; *S. flotator:* one-sample t-test, t = 6.44, d.f. = 21, P < 0.001), and thus leaves effectively amplified the calls. The comparison between both species showed that *H. fleischmanni* calls are amplified to a larger extent than *S. flotator* vocalizations (Two-sample Wilcoxon rank sum test, W = 505, P < 0.001).

For *H. fleischmanni*, in 48% (12/25) of the leaves collected males called from the top, and in the rest (13/25) they called from the underside. The amplitude gains measured for the calls were not affected by the side of the leaf used by males (Two-sample Wilcoxon rank sum test, W = 52, P = 0.277).

### Relationship between leaf dimension, frog position, and call amplification

Not leaf dimensions (i.e., PC1) nor male positions (i.e., PC2) explained the amplitude gain measured for the calls of both species (Table 3, Fig. 5). Male *H. fleischmanni* calls were amplified to a larger extent than *S. flotator* vocalizations, even when accounting for the variation in leaf dimensions or male position between both species (Table 3, Fig. 5).

**Table 3:**
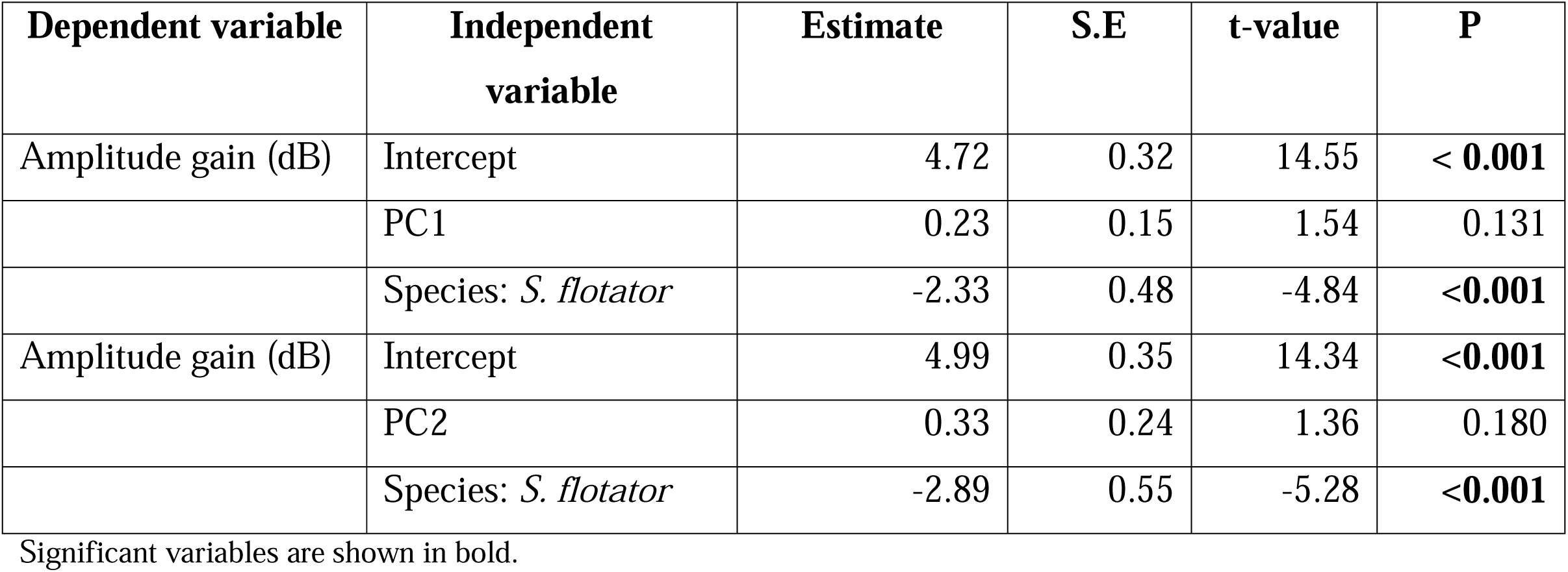
Summary of ANCOVA analyses used to evaluate the effect of leaf dimension and frog position on call amplitude gains.

| **Dependent variable** | **Independent variable** | **Estimate** | **S.E** | **t-value** | **P** |
| --- | --- | --- | --- | --- | --- |
| Amplitude gain (dB) | Intercept | 4.72 | 0.32 | 14.55 | **< 0.001** |
|  | PC1 | 0.23 | 0.15 | 1.54 | 0.131 |
|  | Species: *S. flotator* | -2.33 | 0.48 | -4.84 | **<0.001** |
| Amplitude gain (dB) | Intercept | 4.99 | 0.35 | 14.34 | **<0.001** |
|  | PC2 | 0.33 | 0.24 | 1.36 | 0.180 |
|  | Species: *S. flotator* | -2.89 | 0.55 | -5.28 | **<0.001** |

Significant variables are shown in bold.

**Figure 5:**
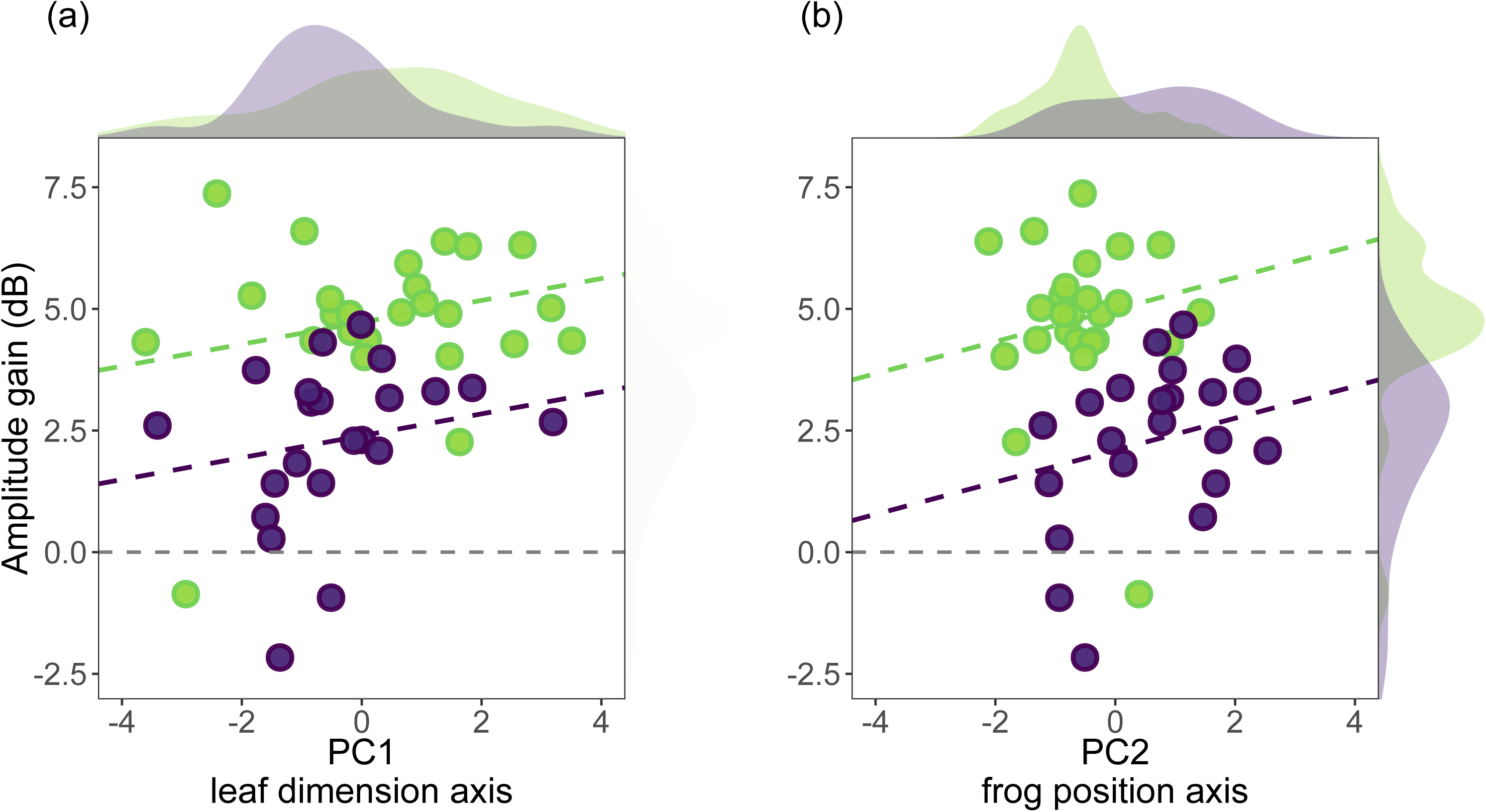
Relationship between call amplitude gain and the scores of (a) PC1 and (b) PC2. Dashed regression lines indicate that slopes were not different from 0. Distribution density plots for each variable are show on the outer margins.

## Discussion

In the present study, we measured the effect of leaf calling sites on the call amplitude of two tropical frogs, an arboreal glass frog and a terrestrial rocket frog species. Overall, there was large overlap between the dimensions of the calling leaves used by both species, although there were some differences between the position of males on their surface. The leaves used by male H. *fleischmanni* amplified the calls of this species by about 5.0 dB, while *S. flotator* leaves moderately amplified their calls by about 2.5 dB. The playback of pure tones showed for the leaves used by both species a frequency band of increased amplification between 4 – 6 kHz. This band matches the frequency content of *H. fleischmanni* calls better than that of *S. flotator’s*. Thus, our results indicate that the different amplitude gains measured for the two species is related to differences in the frequency of their calls, and not in the acoustic properties on the leaves they call from.

Our experimental setup simulated the effect of leaves on the amplitude of calls as perceived by a receiver located above of the calling frog. Because rocket frogs are typically associated to the leaf-litter, amplification above a calling male could facilitate the detection by unintended receiver located above the ground. For example, some birds or parasitic flies could detect these males more easily if their calls are amplified above them.

Unlike other colorful dendrobatids, the brown dorsal coloration of rocket frogs matches well with the surrounding leaf litter, and this species lacks the skin toxins characteristic of other representatives in the family (Mebs et al. 2018). On the contrary, the ventral surface and vocal sac of rocket frogs are lightly colored and conspicuous on the horizontal plane (i.e., when viewed from the forest floor), remaining visually detectable to conspecific males and females. These traits suggest that remaining undetectable to aerial eavesdroppers is advantageous for this terrestrial species, and therefore the leaf amplification effect we measured could impose costs for calling males.

Calling from leaves that amplify more on the horizontal than the vertical plane would favor mate attraction over attracting unintended receivers in terrestrial species. Given that the attenuation of high-frequency sounds can be severe close to the ground (Wiley and Richards 1978), such an effect would be highly beneficial for calling males. Whether leaves also improve sound beaming to the frogs’ horizontal plane is unknown, although there is some evidence of this effect for frogs calling from leaves (Narins and Hurley 1982, Wells & Schwartz 1982) and singing crickets (Erregger and Schmidt 2018). Precisely measuring (e.g., Erregger and Schmidt 2018) or modelling (e.g., Mhatre et al. 2017) the effect of leaves on call radiation along multiple planes would help to elucidate the adaptive role of signaling from these substrates.

Unlike rocket frogs, the arboreal habits of glass frogs indicate that females and unwanted eavesdroppers can be found almost anywhere in the environment around calling males. Frogs calling from the underside will radiate more energy towards the ground, while frogs calling from the top will radiate more energy on the direction of the canopy (Wells and Schwartz 1982, this study). Indeed, calling glass frogs are louder on the leaf side where males were observed calling relative to the opposite side of the leaf (Wells and Schwartz 1982). Although the spatial location of females attending to glass frog choruses is unclear, it has been reported that males calling from higher positions achieve higher reproductive success than males close to the ground (Greer and Wells 1980). This increased success could be explained by favorable above-ground sound propagation (Wiley and Richards 1978), but could also indicate that females approach calling males from the canopy. Therefore, by calling from elevated positions and from the top of leaves male glass frogs could improve detectability by females, although they may be also more exposed to predators and parasites.

Bats are the most likely acoustic eavesdroppers of glass frogs, although katydids are also known to predate on them (Delia et al. 2017). The fringe-lipped bat (*Trachops cirrhosus*) predates on calling frogs, and is attracted by the calls of *H. fleischmanni* (Tuttle and Ryan 1981). Thus, leaf sound reflections that enhance the detection by females may also increase the risk of predation by bats. Male glass frogs calling from the underside of leaves are less exposed to bat attacks. Indeed, geographic variation in the proportion of glass frog calling from the underside is related to the presence of *T. cirrhosus* (Delia et al. 2010). Thus, by flexibly changing the side of the leaf used for calling, male glass frogs can reduce the risk of predation, but also radiate more energy towards the ground, which could decrease the probability of detection by females. Monitoring the effects of males flexibly moving between different sides of calling leaves, and measuring their survival as well as reproductive output would contribute to understand how these frogs balance the attraction of potential mates and predators.

Our results indicate that the spectral differences between the calls of both species explain the degree of amplification they experience, and not the dimensions of the leaf or the position of males. Similar to other vertebrates, the frequency content of frog vocalizations is determined by the size of larynx structures (e.g. Baugh et al. 2018, López et al. 2020). Because larynx size scales with body size, smaller frog species tend to produce higher frequency calls (Tonini et al. 2020). Indeed, male rocket frogs have smaller body size (SVL = 14-17 mm, Savage 2002) than glass frogs (SVL = 19-28 mm, Savage 2002), and consequently call at higher frequencies. Interspecific body size variation in frogs is related to the utilization of different calling posts, such as aquatic and non-aquatic sites (Muñoz et al. 2020), and also to environmental factors relevant for heat and water balance (Amado et al. 2019). For example, male spring pepper frogs (*Pseudacris crucifer*) calling from above the ground are larger and their calls propagate further, but experience a six-fold increase in desiccation rate relative to frogs calling near the ground (Cicchino et al. 2020). Therefore, display sites associated with contrasting environmental conditions can impact body size evolution, and therefore also modify the frequency of vocalizations. Frequency-dependent effects of calling sites are probably more widespread than currently appreciated, and include the leaves investigated here, but also other structures like resonant cavities (e.g, Muñoz and Penna 2016). Furthermore, calling sites may also have a direct impact on sound production mechanisms, with consequences for the frequency content and attractiveness of signals (e.g., Smit et al. 2019, Goutte et al. 2020, Muñoz et al. 2020). Yet, understanding how evolutionary changes in intrinsic factors, such as body size, can impact the way organism interact with display sites is still a gap in our knowledge.

The scale at which animals interact with their environment for effective communication needs to be investigated at many different levels. Here we show that the leaves used by a terrestrial and an arboreal frog species amplify their calls, but argue that this effect needs to be considered in relation to the broad-scale environment and also the ecological singularities of different species. Animals can display from diverse sites in the environment, yet they often select specific locations, and the adaptive significance of these choices are just beginning to be understood.

## Notes

**<u>Funding:</u>** This work was supported by Becas Chile 2018-Comisión Nacional de Investigación Científica y Tecnológica (CONICYT) scholarship awarded to MIM (number: 72190501).

### Competing Interest Statement

The authors have declared no competing interest.

